## Supplementary material for "Robust detection of oncometabolic aberrations by ^1^H-^13^C heteronuclear single quantum correlation in live cells and intact tumors *ex-vivo*": Barekatain et.al Supplemental Info

**Supplementary Figure S1: 2-HG is dramatically elevated in mutant *IDH1* cells but the detection of 2-HG specific peaks is unreliable in the $^{1}H$ spectrum due to extensively convolution by signals from high abundant metabolites.**

**Supplementary Figure S2: Signal to noise ratio (SNR) analysis of $^{1}H$ projections from the HSQC spectrum of individual peaks in 2-HG chemical standard.**

**Supplementary Figure S3: H-C3-H peaks of 2-HG are uniquely detected in HSQC spectra *IDH1*-mutant cells and are eliminated by mutant IDH1 specific inhibitor treatment.**

**Supplementary Figure S4: Gluconate and 6-PG are highly abundant metabolites in *PGD*-deleted cells but are indistinct in the $^{1}H$ spectrum of *PGD*-deleted tumors because of nearness of the water signal and spectral convolution by high abundant metabolites.**

**Supplementary Figure S5: Phase sensitive $^{1}H$-$^{13}C$ HSQC spectrum of gluconate standard with 1H projection.**

**Supplementary Figure S6: Specific detection of gluconate in *PGD*-deleted but not PGD-rescued or WT tumors.**

#### Supplementary Figure S1

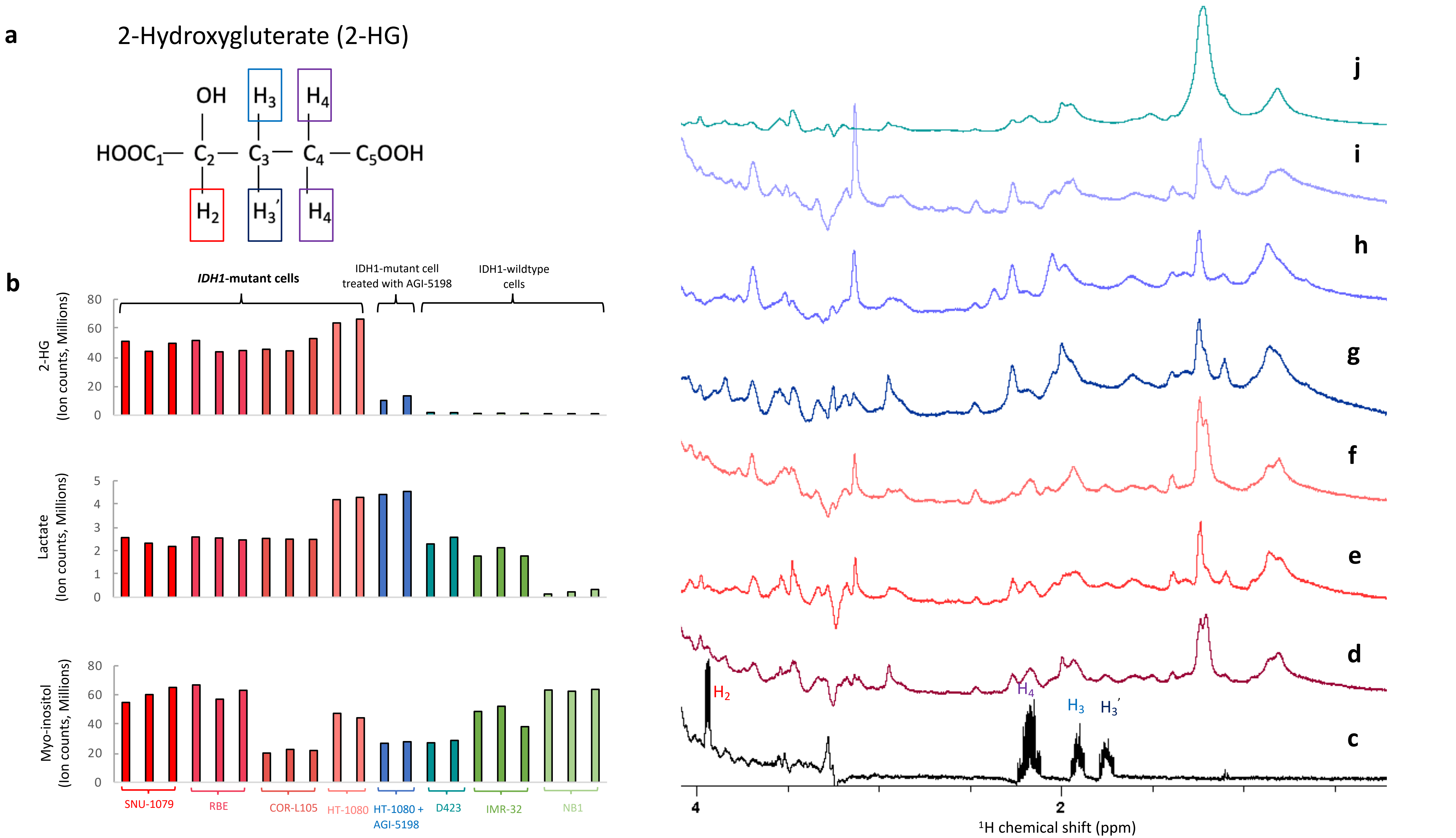

**Supplementary Figure S1: 2-HG is dramatically elevated in mutant *IDH1* cells but the detection of 2-HG specific peaks is unreliable in the $^{1}$H spectrum due to extensively convolution by signals from high abundant metabolites.** (**a**) The molecular structure of 2-HG. (**b**) The mass spectroscopy data comparing the level of 2-HG, lactate and Myo-inositol in IDH1 mutant cells (HT-1080 and NHA-IDH1) and wildtype cells (NHA and D423). The average value of 2-HG in IDH1 mutant cells is (51+/-7) *10^6 ion counts while this value for HT-1080 cell treated with inhibitor is (11+/-1.6)*10^6 ion counts and for IDH1 wildtype cells is (1+/-.5)*10^6 ion counts. While the level of lactate and myo-inositol do not vary significantly between IDH1 and wildtype cells. Based on mass-spect data, 2-HG is high abundant metabolites in IDH1 mutant cells therefore it can be detected by magnetic resonance spectroscopy. $^{1}$H spectrum of (**c**) 2-HG standard and its peaks assignment, (**d**) live R132C IDH1 mutant cells (HT-1080), (**e**) live R132H IDH1 mutant cells (NHA IDH1), (**f**) live R132C IDH1 mutant cells (SNU1079), (**g**) live R132C IDH1 mutant cells (HT-1080) treated with 10uM mutant IDH1 inhibitor (AGI-5198) for 48hrs (**h**) live R132H IDH1 mutant cells (NHA IDH1) treated with 10uM mutant IDH1 inhibitor (AGI-5198) for 48hrs (**i**) live R132C IDH1 mutant cells (SNU-1079) treated with 10uM mutant IDH1 inhibitor (AGI-5198) for 48hrs (**j**) live IDH1 wild type cells (D423). Due to the peaks broadening and presence of other highly abundant metabolites, 1H spectrum does not allow reliable detection of 2-HG peaks.

#### Supplementary Figure S2

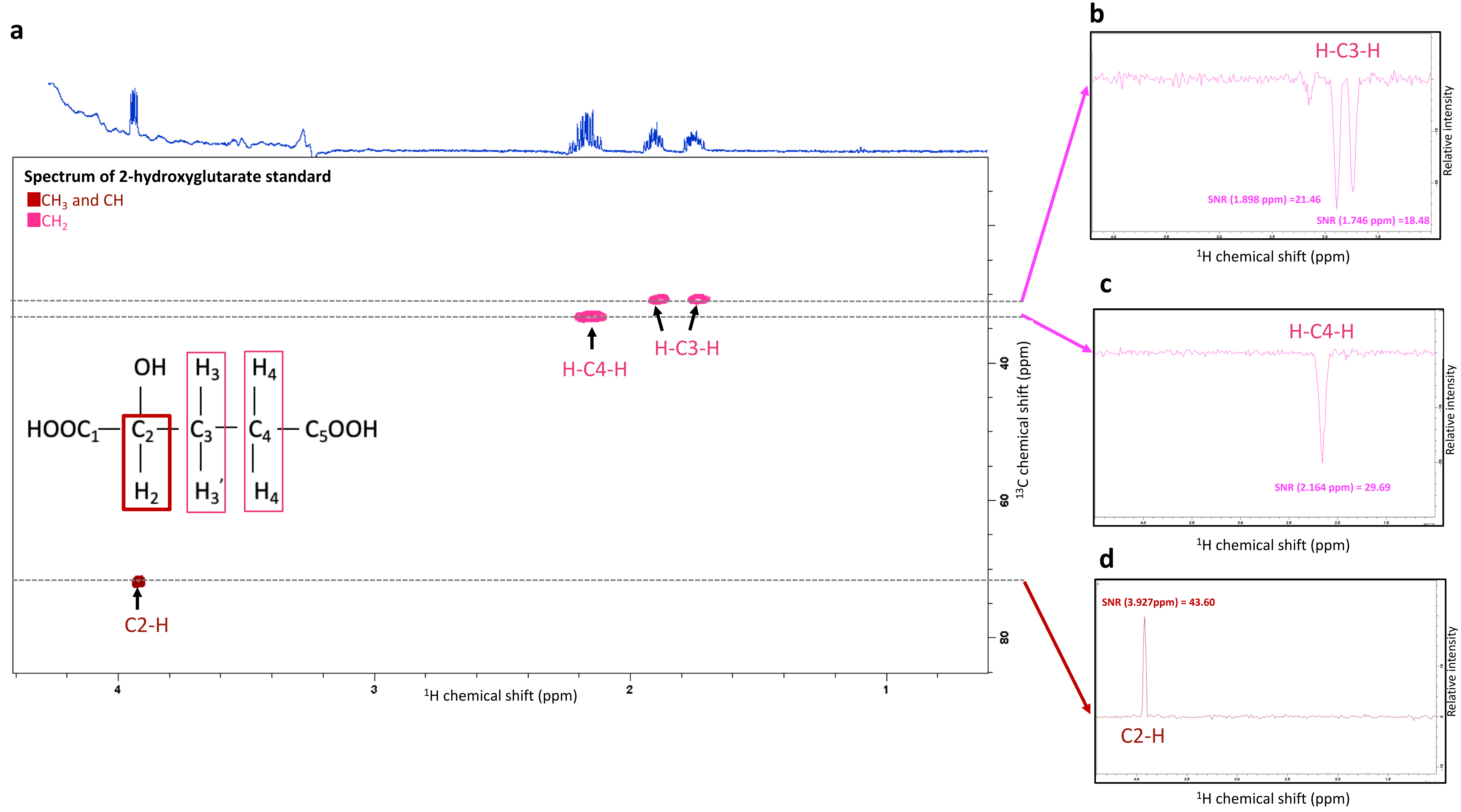

**Supplementary Figure S2: Signal to noise ratio (SNR) analysis of $^{1}H$ projections from the HSQC spectrum of individual peaks in 2-HG chemical standard.** Phase sensitive $^{1}H$-$^{13}C$ HSQC spectrum of a chemical standard 2-HG at 7.5mM concentration in 90% PBS and 10% $D_2O$ using the HSQCEDETGPSISP2.3 pulse sequence, where the first color code in each spectrum depicts peaks which have positive phase ($CH_3$ and CH) and the second color code shows peaks which have negative phase ($CH_2$). The spectrum is acquired using 500 MHz Bruker AVANCE III NMR. The 1D $^{1}H$ spectrum of 2-HG standard displayed as the x-axis on top of the 2D HSQC spectrum. In $^{1}H$-$^{13}C$ HSQC, the protons which are directly bounded with $^{13}C$ atoms are detected, these carbons and protons are labeled in the 2-HG structure, and peaks are also annotated in the spectrum. The C2-H peak in the phase sensitive $^{1}H$-$^{13}C$ HSQC spectrum has the positive phase, while the other peaks have negative phase (H-C-H). Phases of each peak can be more appreciated from their $^{1}H$ projection of the row containing each peak as shown on the right side of the figure. The signal to noise ratio for each peak is acquired by adjusting the phase of negative peaks to positive and selecting the region of interest and applying SINO command in Topspin. The small bump in the 1D projection of H-C3-H peak is caused by the impurity in our standard.

### Supplementary Figure S3

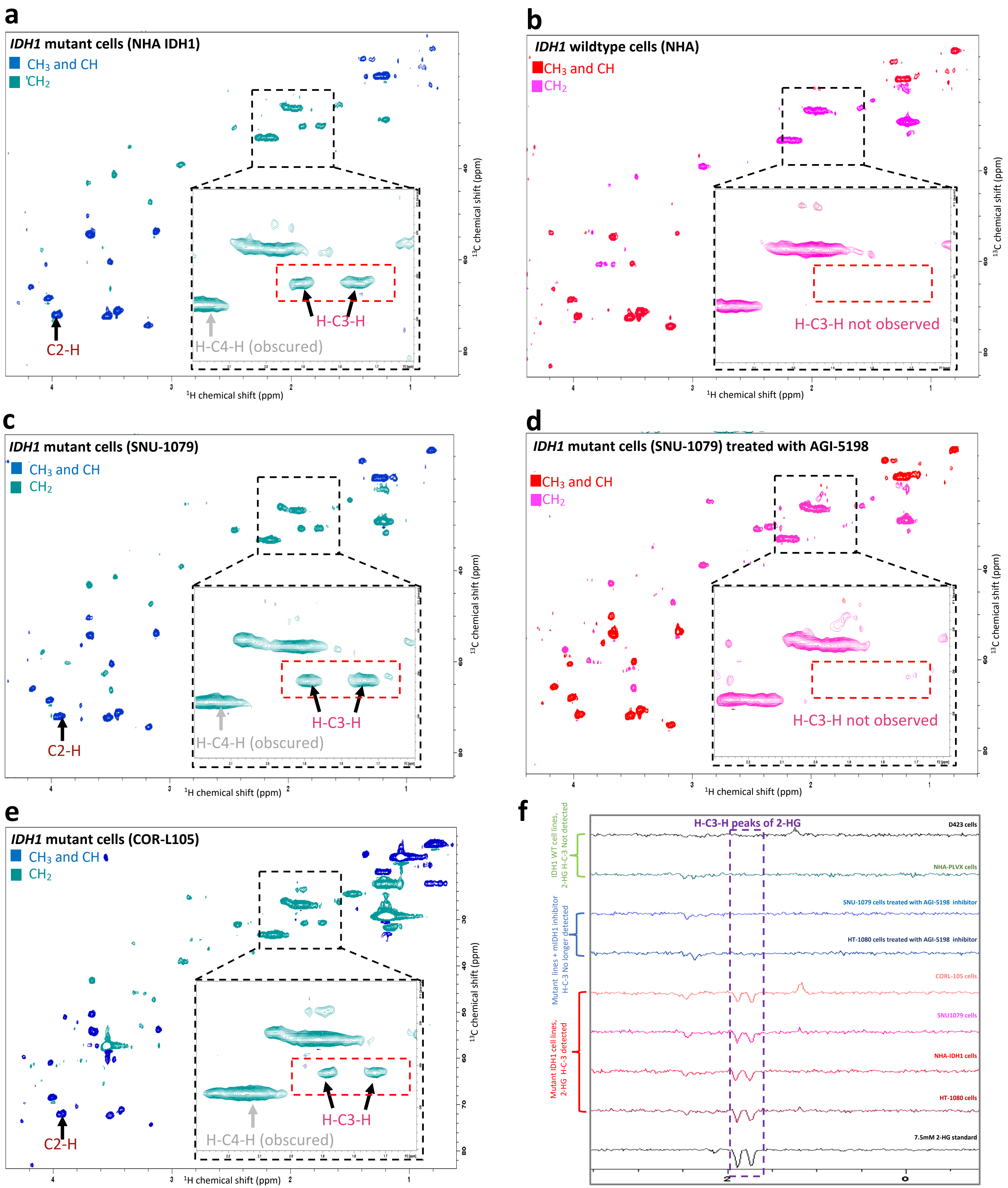

**Supplementary Figure S3: H-C3-H peaks of 2-HG are uniquely detected in HSQC spectra *IDH1*-mutant cells and are eliminated by mutant IDH1 specific inhibitor treatment.** Phase sensitive $^{1}H$-$^{13}C$ HSQC spectra using HSQCEDETGPSISP2.3 pulse sequence, where the first color code in each spectrum depicts the peaks which have positive phase ($CH_3$ and CH) and the second color code shows the peaks which have negative phase ($CH_2$), acquired using 500 MHz Bruker AVANCE III NMR. The red dashed box shows where we expect to see H-C3-H peaks of 2-HG in the spectrum. (**a**) The spectrum of live IDH1 mutant cells (immortalized normal human astrocyte, NHA IDH1) cells which express mutant IDH1. (**b**) The $^{1}H$-$^{13}C$ HSQC spectrum of live immortalized normal human astrocyte cells (NHA) which express non-mutant IDH1. (**c**) The $^{1}H$-$^{13}C$ HSQC spectrum of IDH1 mutant cells (SNU-1079). (**d**) The $^{1}H$-$^{13}C$ HSQC spectrum of IDH1 mutant cells (SNU-1079) treated with 10uM mutant IDH1 inhibitor (AGI-5198) for 48hrs. (**e**) The spectrum of live IDH1 mutant cells (CORL-105). The presence of 2-HG peaks is readily evident in spectra of all IDH1 mutant cells (**a**,**c** and **e**)**.** C2-H peak of 2-HG has the chemical shift close to the highly abundant myo-inositol peak, and barely distinguishable. The H-C4-H peak of 2-HG is obscured by the broad (-CH2-) lipid peak. However, H-C3-H peaks of 2-HG are easily detectable in the IDH1 mutant cells spectrum. The spectrum of live IDH1 wildtype cells (NHA) shows complete absence of H-C3-H peaks of 2-HG **(b).** Also, treating cells with mutant IDH1 inhibitor results in disappearance of H-C3-H peaks of 2-HG (**d**)**.** (**f**) Shows the $^{1}H$ projection of H-C3-H peaks of 2-HG from $^{1}H$-$^{13}C$ HSQC spectrum of 2-HG standard, live mutant IDH1 cells, live mutant IDH1 cells treated with mutant IDH1 inhibitor and live IDH1 wildtype cells. $^{1}H$ projection extracted from 2D HSQC spectrum of mutant IDH1 cells show the negatively phased doublet associated with H-C3-H peaks of 2-HG. The spectra are normalized to the noise level.

#### Supplementary Figure S4

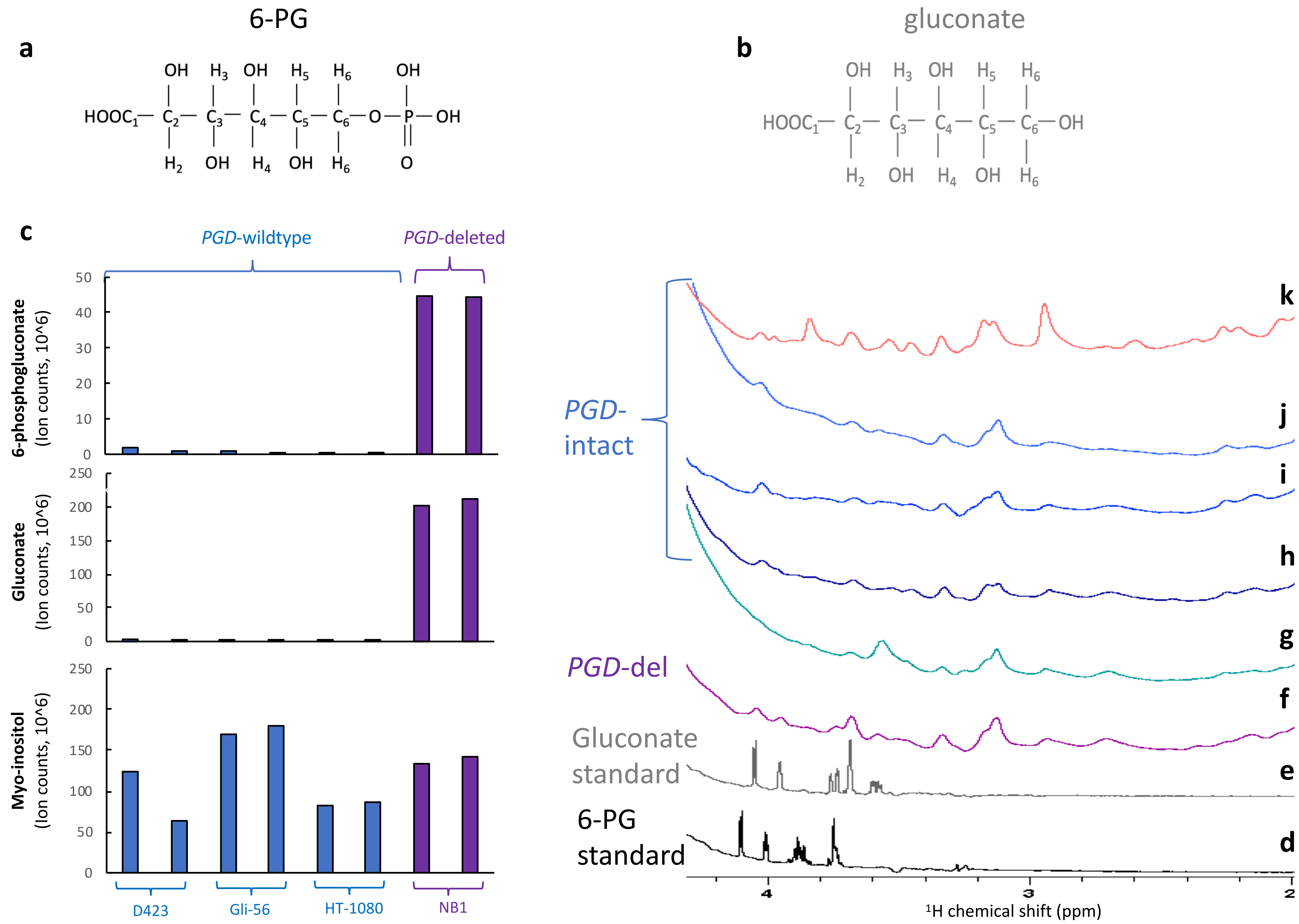

**Supplementary Figure S4: Gluconate and 6-PG are highly abundant metabolites in *PGD*-deleted cells but are indistinct in the $^{1}H$ spectrum of *PGD*-deleted tumors because of nearness of the water signal and spectral convolution by high abundant metabolites:** (**a**) The molecular structures of gluconate and (**b**) 6-phosphogluconate. (**c**) mass-spec data comparing the levels of 6-PG, gluconate and lactate in extracts of *PGD*-deleted vs wildtype cells indicates that *PGD*-deletion leads to a dramatic accumulation of 6-PG/gluconate. Yet, 6-PG or gluconate are not readily identifiable and do not distinguish the $^{1}H$ spectra of *PGD*-deleted xengrafted tumors *ex-vivo* from those that are *PGD*-intact or rescued. $^{1}H$ spectrum of (**d**) 9 mM 6-PG standard and (**e**) 9 mM gluconate standard in 90% PBS and 10% $D_2O$ (**f**) *PGD*-deleted tumor (NB1) and (**g**) *PGD*-rescued tumor (NB1-*PGD*) (**h**) *PGD*-wildtype tumors (D423) (**i**) *PGD*-wildtype tumor (G59) (**j**) *PGD*-wildtype tumor (U87) (**k**) normal mouse brain. Inability to visualized 6-PG/gluconate in PGD-deleted tumors is most likely due to spectral convolution by chemical shifts from other highly abundant metabolites and closeness of gluconate chemical shifts to the water signal.

#### Supplementary Figure S5

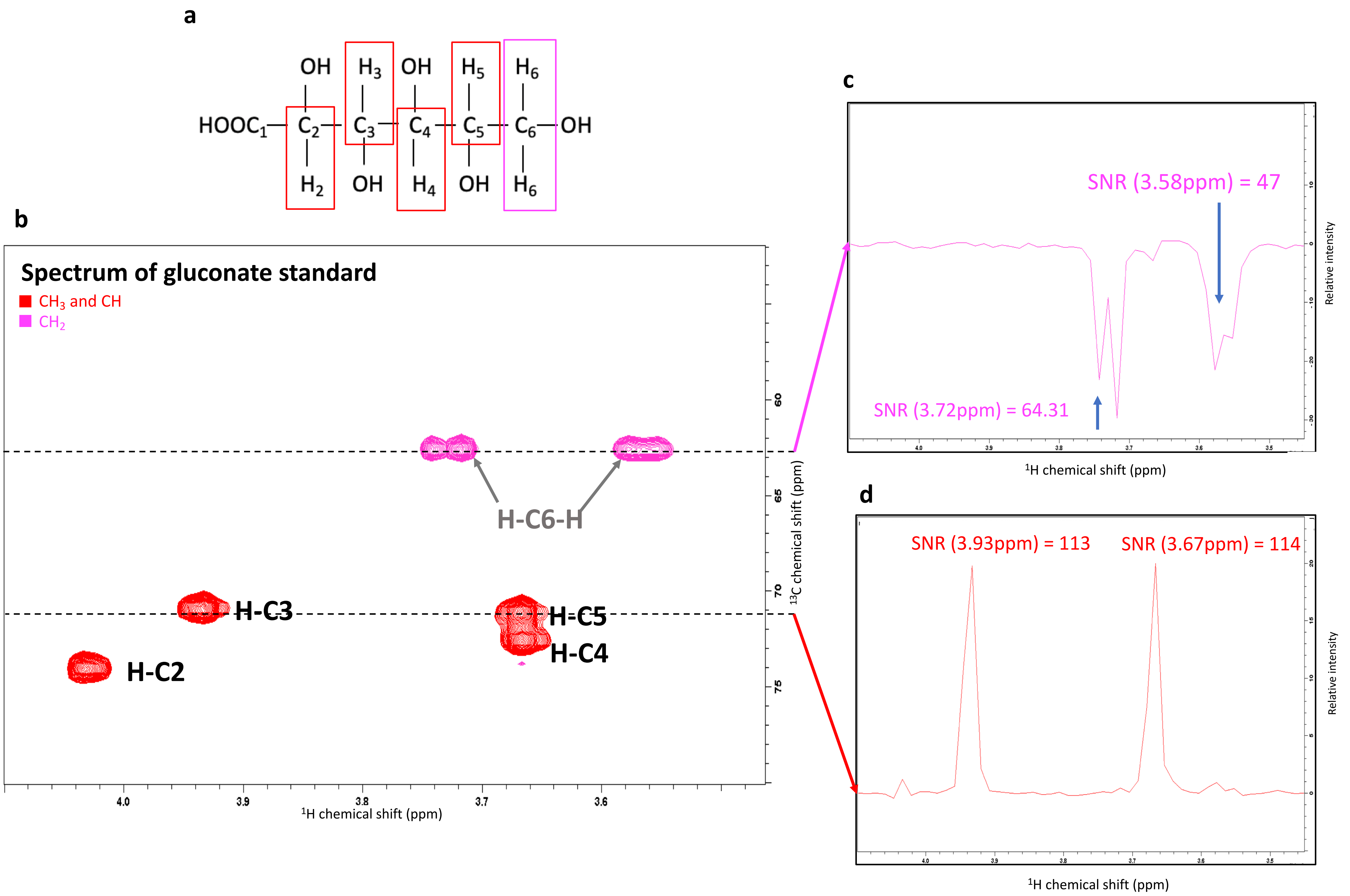

**Supplementary Figure S5: Phase sensitive $^{1}H$-$^{13}C$ HSQC spectrum of gluconate standard with 1H projection.** (**a**) Molecular structure of gluconate. (**b)** The Phase sensitive $^{1}H$-$^{13}C$ HSQC spectrum of 9 mM gluconate standard in PBS and 10% $D_2O$ using HSQCEDETGPSISP2.3 pulse sequence. Red peaks are associated with C-H groups of gluconate and have positive phase while purple peaks are associated with H-C-H group of gluconate and have negative phase. (**c** and **d**) show the $^{1}H$ projection of rows containing H-C6-H and H-C5 and H-C3 peaks of gluconate with signal to noise ratio associated from each peak. For the sake of simplicity, of the figure, the $^{1}H$ projection of other peaks are now shown here. For the H-C4 peak, the SNR(3.67 ppm)= 126 and for the H-C4 peak SNR(4.04 ppm)=157.

#### Supplementary Figure S6

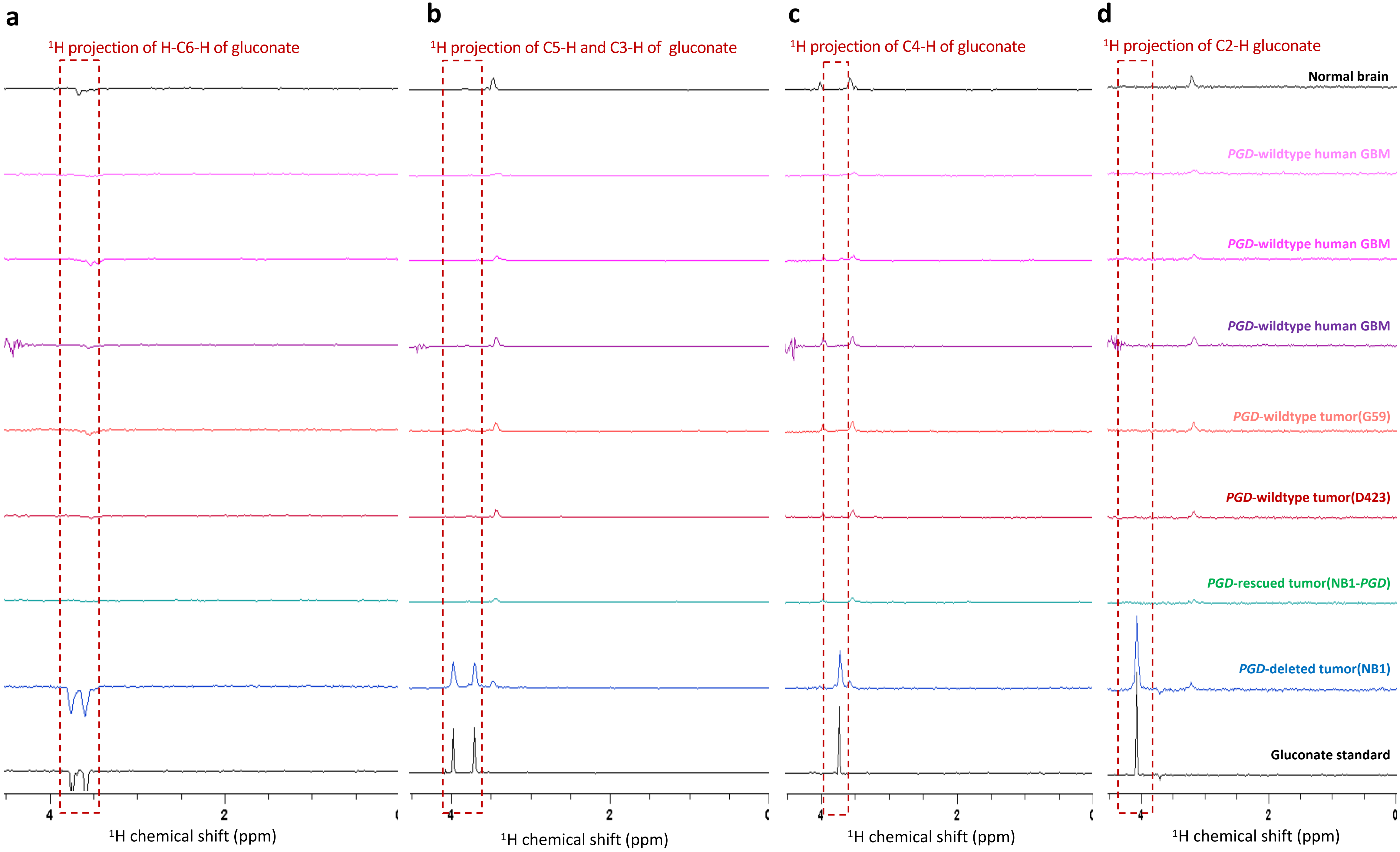

**Supplementary Figure S6: Specific detection of gluconate in *PGD*-deleted but not PGD-rescued or WT tumors.** The $^{1}H$ projections of rows containing (**a**) H-C6-H peaks (**b**) C5-H and C3-H (**c**) C4-H and (**d**) C2-H of gluconate from the phased sensitive $^{1}H$-$^{13}C$ HSQC spectrum of 9 mM gluconate standard, *PGD*-deleted tumors, *PGD*-rescued tumors, *PGD* wildtype tumors( xenografted and human GBM) and normal mouse brain. $^{1}H$ projections extracted from 2D HSQC spectrum of *PGD*-deleted tumors show the peaks associated with gluconate peaks. While these peaks are absent in *PGD*-rescued and wildtype tumors and normal mouse brain. The spectra are normalized to the noise level.
